# Cell-type-specific synaptic imbalance and disrupted homeostatic plasticity in cortical circuits of ASD-associated *Chd8* haploinsufficient mice

**DOI:** 10.1101/2020.05.14.093187

**Authors:** Robert A. Ellingford, Emilie Rabeshala de Meritens, Raghav Shaunak, Liam Naybour, M. Albert Basson, Laura C. Andreae

**Affiliations:** Centre for Developmental Neurobiology, Institute of Psychiatry, Psychology & Neuroscience, King’s College London, UK; Centre for Craniofacial & Regenerative Biology, King’s College London, UK; MRC Centre for Neurodevelopmental Disorders, King’s College London, UK

## Abstract

Heterozygous mutation of chromodomain helicase DNA binding protein 8 (*CHD8*) is strongly associated with autism spectrum disorder (ASD) and results in dysregulated expression of neurodevelopmental and synaptic genes during brain development. To reveal how these changes affect ASD-associated cortical circuits, we studied synaptic transmission in the prefrontal cortex of a haploinsufficient *Chd8* mouse model. We report profound alterations to both excitatory and inhibitory synaptic transmission onto deep layer projection neurons, resulting in a reduced excitatory:inhibitory balance, which were found to vary dynamically across neurodevelopment and result from distinct effects of reduced *Chd8* expression within individual neuronal subtypes. These changes were associated with disrupted regulation of homeostatic plasticity mechanisms operating via spontaneous neurotransmission. These findings therefore directly implicate *CHD8* mutation in the disruption of ASD-relevant circuits in the cortex.

## Introduction

Autism spectrum disorder (ASD) is a common neurodevelopmental disorder characterized by social communication deficits and repetitive behaviours. Imbalances in the levels of excitatory and inhibitory activity (E:I balance) within the neuronal circuitry of the cerebral cortex are a proposed causative mechanism of ASD (1–3). E:I balance governs the output of neurons based on the local level of excitatory/inhibitory synaptic transmission and intrinsic excitability. Some of the highest confidence ASD-risk genes encode proteins that directly impact E:I balance (4,5) and multiple animal models display synaptic transmission deficits alongside ASD-like behavioral abnormalities (2,6–12).

E:I balance is sustained through homeostatic plasticity mechanisms that maintain network activity within an optimal range by tuning synaptic strength (13) and intrinsic excitability (14). However, the role that homeostatic plasticity plays in the context of ASD-risk mutations remains unclear. It has been proposed that homeostatic mechanisms could be insufficient or maladaptive (2). Reductions in normal homeostatic responses have been demonstrated in dissociated neuronal cultures from mice with mutations in the ASD-associated genes *MeCP2* (15–17), *Fmr1* (18,19) and *Shank3* (20) and in visual cortex following eyelid suture (20), implicating the failure of such mechanisms in ASD etiology. Conversely it has been suggested that many changes to E:I balance seen in ASD gene mutant mice may themselves be compensatory and reflect the normal operation of homeostatic plasticity (21). Notably, there has been a lack of study in regions of the brain directly relevant to ASD, such as prefrontal cortex (PFC). Furthermore, to what extent changes to synaptic E:I balance are specific to the cell type affected by the gene deletion versus secondary or even compensatory effects remains poorly understood.

*De novo* heterozygous mutation of chromodomain helicase DNA binding protein 8 (*CHD8*) is one of the highest confidence genetic risk factors for ASD (22–26). *CHD8* encodes a chromatin remodeller that regulates the expression of many other ASD-associated genes essential for brain and synapse development in both human (27–30) and mouse models (7,8,31–33). However, despite these studies, the link between reduced *CHD8* expression and a definitive pathological mechanism that can give rise to ASD remains unclear. We therefore conducted a detailed characterization of E:I balance during postnatal cortical development in a *Chd8* heterozygous (*Chd8*^+*/-*^) mouse model (32). We focused our analysis on deep layer (V/VI) pyramidal neurons within the PFC as this cell type and brain region have been strongly implicated in ASD etiology through both anatomical (34) and transcriptomic (35) studies. These mice were found to display developmental stage-specific alterations to both excitatory and inhibitory synaptic transmission. *Chd8* haploinsufficiency targeted to excitatory or inhibitory neurons revealed both cell-type-specific synaptic alterations and indirect changes. Finally, we found that homeostatic responses, dependent on spontaneous neurotransmission, were defective in the *Chd8*^+*/-*^ PFC.

## Materials and Methods

### Mice

All procedures were performed according to the Animals (Scientific Procedures) Act 1986 with ethical approval granted by the UK Home Office. The conditional *Chd8* (*Chd8*^*flox*^) and *Chd8* null (*Chd8*^*-*^) alleles were generated by our lab and have been described in detail previously (32). *Chd8*^*flox*^ mice were bred with a variety of *Cre* mice to excise exon 3, resulting in an early frameshift and termination of translation at amino acid 419 (*Chd8*^*-*^) to produce a protein lacking all functional domains, equivalent to nonsense and frameshift mutations identified in patients (36). Constitutive *Chd8* heterozygotes (*Chd8*^+*/-*^) have previously been verified to express approximately 50% of the WT levels of *Chd8* at both the RNA and protein level (32).

For dendritic spine and inhibitory synapse analysis, *Chd8*^+*/-*^ mice were bred with Tg(Thy1-EGFP)MJrs/J (*Thy1-GFP-M*) mice (37). For experiments utilising conditional *Chd8*^+*/-*^ mice (*cChd8*^+*/-*^), homozygous *Chd8*^*flox/flox*^ mice were bred with mice heterozygous for the respective Cre line (*Nkx2.1-Cre* (38) *or NEX-Cre* (39)). All genotyping used the HotSHOT method for DNA extraction (40).

### Electrophysiology

Electrophysiological recordings were performed on acute brain slices using whole-cell patch clamp electrophysiology. For details of standard electrophysiological methods: slice preparation (including use of Na^+^ reintroduction (41) for adult slices), miniature postsynaptic current and intrinsic properties recordings, please see Supplementary Materials and Methods.

#### Homeostatic plasticity recordings

P13-15 mouse brains were sliced as detailed in Supplementary materials and methods. Before being placed in ACSF to recover, slices were cut down the midline in order to separate the two hemispheres. One hemisphere was placed in standard ACSF and the other placed in ACSF containing drug treatment (1 µM TTX + 25 µM D-APV, 1 µM TTX + 10 µM GZ or 1 µM TTX alone). Slices were incubated for 6 hours at room temperature with perfusion of 95% O_2_/5% CO_2_ before drug treated slices were transferred to standard ACSF. Miniature post synaptic currents were then recorded from the treated and untreated slices as before. To allow comparison between animals, the frequency and amplitude of currents for each animal were normalized to the mean untreated value.

### Sholl Analysis

For details of dendritic imaging and Sholl analysis, please see Supplementary Materials and Methods.

### Synapse Analysis

For details of immunohistochemistry, image acquisition and analysis, please see Supplementary Materials and Methods

### Statistics

All data is presented as mean ± standard error of the mean (SEM) unless otherwise stated. All statistical analyses were performed using Prism software (Graphpad). For pairwise comparisons the D’Agostino-Pearson omnibus normality test was first used to determine Gaussian distribution within datasets. If both data sets were normally distributed, comparisons were performed using unpaired t tests (with Welch’s correction if standard deviations were found to significantly differ). If data was not normally distributed, then pairwise comparisons were performed using the nonparametric Mann-Whitney U test (Supplementary Table 1). Multivariate comparisons were performed using a two-way ANOVA followed by Tukey’s multiple comparisons test (Supplementary Table 2).

## Results

### Reduced synaptic E:I balance in the PFC of Chd8^+/-^ mice

We first determined the effect of *Chd8* haploinsufficiency on the balance of synaptic transmission within the PFC. Whole-cell voltage clamp recordings of miniature excitatory and inhibitory postsynaptic currents (mEPSCs and mIPSCs respectively) were performed in deep layer neurons of the PFC, using *ex vivo* brain slices prepared from postnatal day 20 (P20) *Chd8*^+*/-*^ mice and their wild-type (WT) littermates. *Chd8*^+*/-*^ neurons displayed significantly decreased frequency and amplitude of mEPSCs (Fig. 1a) and increased mIPSC frequency (Fig. 1b), indicating decreased levels of excitatory synaptic transmission with a concurrent increase in inhibitory transmission. We then determined the intrinsic excitability of these same neurons by measuring the frequency of action potential (AP) firing in response to increasing current injection (*f-I* curves) at P19-P21 using whole-cell current clamp recordings. The *f-I* curves of WT and *Chd8*^+*/-*^ neurons did not differ significantly (Fig. 1c). In addition, the magnitude and width of representative APs were equivalent between genotypes, as well as the voltage threshold and rheobase required to elicit responses (Fig. 1d). We also compared the passive membrane properties of WT and *Chd8*^+*/-*^ neurons, finding no differences in resting membrane potential (RMP), membrane capacitance or membrane resistance (Fig. 1e). Therefore, reduced *Chd8* expression does not affect neuronal intrinsic excitability, size or ion channel composition. Finally, we detected no large-scale changes to neuronal structure as Sholl analysis revealed there to be no alterations in the degree of dendritic arborization in P22 Golgi-Cox stained *Chd8*^+*/-*^ neurons (Fig. 1f). Together these results suggest that reduced *Chd8* expression specifically impacts the development of synapses. Coupled with unaltered intrinsic excitability, the observed changes to synaptic transmission in *Chd8*^+*/-*^ neurons will result in reduced E:I balance within the PFC compared to WT littermates.

**Fig. 1:**
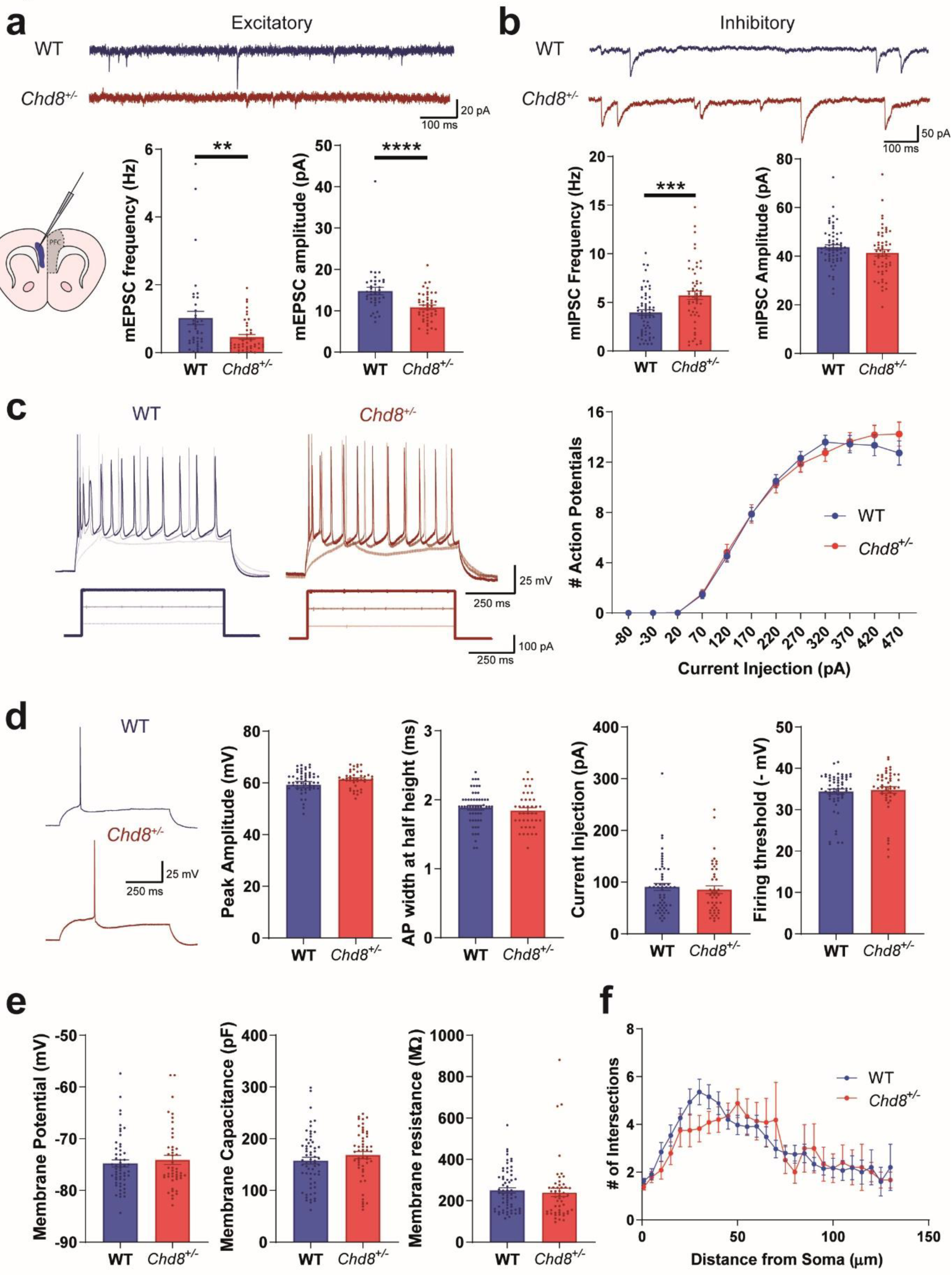
Reduced synaptic E:I balance in the PFC of Chd8^+/-^ mice. (**a, b**) Representative mEPSC (**a**) and mIPSC (**b**) recordings from pyramidal neurons within layer V/VI of the PFC (blue area on schematic indicates recording area) of *ex vivo* slices from P19-21 WT (*blue*) and *Chd8*^*+/-*^ (*red*) animals alongside quantifications of mEPSC and mIPSC frequency (*bottom left graphs*) and amplitude (*bottom right graphs*). (**a**) *Chd8*^*+/-*^ neurons display significantly reduced mEPSC frequency (*p*=0.0027) and amplitude (*p*<0.0001) but (**b**) increased mIPSC frequency (*p*=0.0009) and equivalent mIPSC amplitude. (**c**) Normal action potential (AP) firing frequency in *Chd8*^*+/-*^ neurons; *left:* representative voltage traces and current injection stimuli (*below*), *right: f-I* curve. (**d**) Representative individually-elicited APs. *Chd8*^*+/-*^ neurons showed no difference in AP amplitude (*left*), width (*middle left*), rheobase (*middle right*), or firing threshold (*right*). (**e**) Passive membrane properties: *Chd8*^*+/-*^ neurons showed no difference in resting membrane potential (*left*), whole-cell capacitance *(middle*) or membrane resistance (*right*). (**f**) Sholl analysis of basal dendrites (P22) shows no significant difference in *Chd8*^*+/-*^ neurons. For all Fig.s, bars show mean, error bars SEM, **p<0.01, ***p<0.001, ****p<0.0001.

### Developmental stage-specific synaptic effects of Chd8 heterozygosity

*Chd8* expression levels and the effects of *Chd8* haploinsufficiency on gene expression vary substantially over the course of neurodevelopment (31,32). Therefore, in addition to P20 animals, we characterised excitatory and inhibitory synaptic integrity within *Chd8*^+*/-*^ mice over a time course consisting of neonatal (P5), adolescent (P14) and adult (P55-60) ages. We found no significant differences in mEPSC frequency until the decrease at the P20 timepoint which appeared to persist into adulthood (Fig. 2a). We also observed that mEPSC amplitude was already decreased at P14 but had recovered by adulthood (Fig. 2a). We saw no alterations to mIPSC frequency apart from at P20 but did observe substantial alterations to mIPSC amplitude, which was decreased at P5 and increased at P14 and adult stages (Fig. 2b). To determine whether these alterations in frequency resulted from structural changes to synapse number we assessed the density of dendritic spines and vesicular GABA transporter (VGAT)-positive synapses on the secondary apical and basal dendrites of these neurons at P14 and P20 [see Supplementary methods]. We saw no difference in dendritic spine or VGAT+ synaptic density on apical or basal dendrites at P14 (Fig. 2c), consistent with unaltered mEPSC and mIPSC frequency at this time point. In P20 neurons we saw no change in dendritic spine density but did see an increased density of VGAT+ synapses specifically on basal dendrites (Fig. 2d). Therefore, the decrease in mEPSC frequency at P20 is not caused by reduced excitatory synaptic density per se, but more likely by functional changes such as decreased glutamate release probability. In contrast, the increased mIPSC frequency observed at this time point can at least partially be ascribed to an increase in the number of inhibitory synapses onto basal dendrites.

**Fig. 2:**
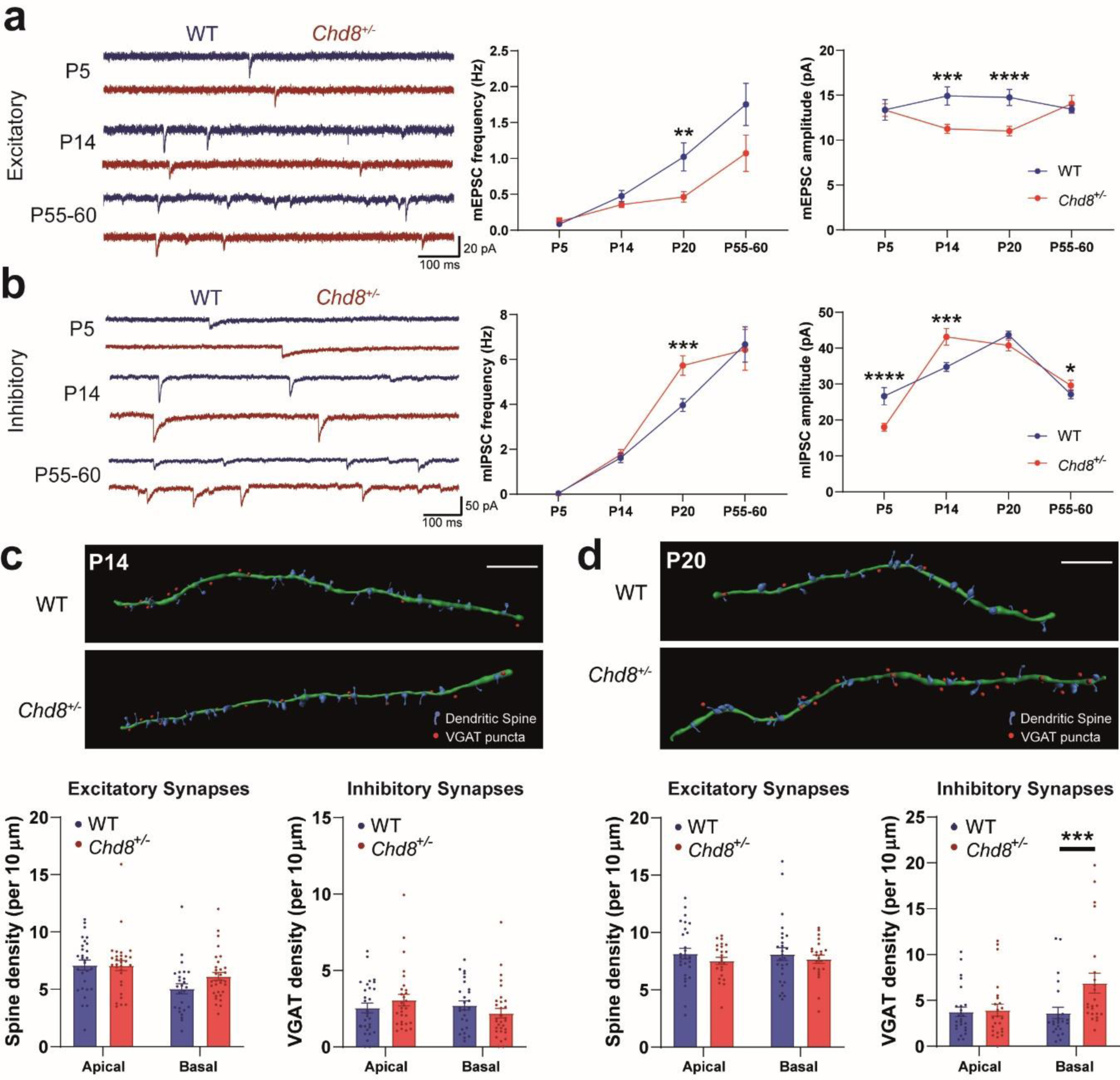
Developmental stage-specific synaptic effects of Chd8 heterozygosity. (**a, b**) Developmental trajectory of excitatory (**a**) and inhibitory (**b**) synaptic function; *left:* representative mEPSC (**a**) and mIPSC (**b**) recordings from PFC neurons in *ex vivo* slices from P5, P14 and P55-60 WT (*blue*) and *Chd8*^*+/-*^ (*red*) animals: frequency (*middle*) and amplitude (*right*) changes during development. *Chd8*^*+/-*^ neurons showed no significant difference in mEPSC frequency at P5, P14 and P55-60 nor in amplitude at P5 and P55-60 but did show decreased mEPSC amplitude at P14. *Chd8*^*+/-*^ neurons showed no significant difference in mIPSC frequency at P5, P14 and P55-60, but displayed decreased mIPSC amplitude at P5 (*p*<0.0001) followed by increased amplitude at P14 (*p*=0.0008) and P55-60 (*p*=0.033). (**c, d**) Analysis of synapse densities at P14 (**c**) and P20 (**d**); top: representative reconstructions of secondary basal dendrites (*green filament*) with associated spines (*blue*) and inhibitory synapses (*red dots*) from WT (*above*) and *Chd8*^*+/-*^ (*below*) *GFP+* pyramidal neurons immunolabelled for the inhibitory synapse marker VGAT, with quantifications of spine (*bottom left graphs*) and VGAT (*bottom right graphs*) densities (scale bar 10 µm). At P14, *Chd8*^*+/-*^ neurons showed no difference in either spine or inhibitory synapse densities. At P20, *Chd8*^*+/-*^ neurons showed no difference in spine densities, nor in VGAT density on apical dendrites but did show an increase in VGAT density on basal dendrites (*p*=0.0002). *p<0.05, **p<0.01, ***p<0.001, ****p<0.0001.

### Differential alterations to synaptic transmission from cell-type-specific Chd8 heterozygosity

To determine whether these contrasting synaptic phenotypes arise through cell autonomous effects of reduced *Chd8* expression we selectively targeted either excitatory neurons using a forebrain-specific, post-mitotic Cre driver crossed with a conditional *Chd8*^*flox*^ allele (*NEX-cChd8*^+*/-*^), or the majority of inhibitory neurons using the *Nkx2.1-Cre* driver (*Nkx2.1-cChd8*^+*/-*^) and performed mEPSC and mIPSC recordings in the PFC as before, at P19-21. Haploinsufficiency of *Chd8* solely in interneurons (*Nkx2.1-cChd8*^+*/-*^) replicated the increase in mIPSC frequency seen in constitutive *Chd8* heterozygotes while exhibiting no changes to excitatory transmission (Fig. 3a), indicating that the inhibitory transmission phenotype is driven cell-autonomously by interneurons. Equally, *Chd8* haploinsufficiency restricted to excitatory neurons (*NEX-cChd8*^+*/-*^) replicated the decreased mEPSC frequency seen in full *Chd8* heterozygotes (Fig. 3b), suggesting that *Chd8* regulates excitatory transmission cell-autonomously in post-mitotic excitatory neurons. However, in contrast to the constitutive *Chd8* heterozygotes, *NEX-cChd8*^+*/-*^ PFC neurons also displayed increased mEPSC amplitudes and decreased mIPSC frequency (Fig. 3b), suggestive of activation of compensatory mechanisms. While these are likely to be complex, the absence of such compensation in the constitutive *Chd8*^+*/-*^ mice led us to hypothesize that homeostatic mechanisms might be impaired in these animals.

**Fig. 3:**
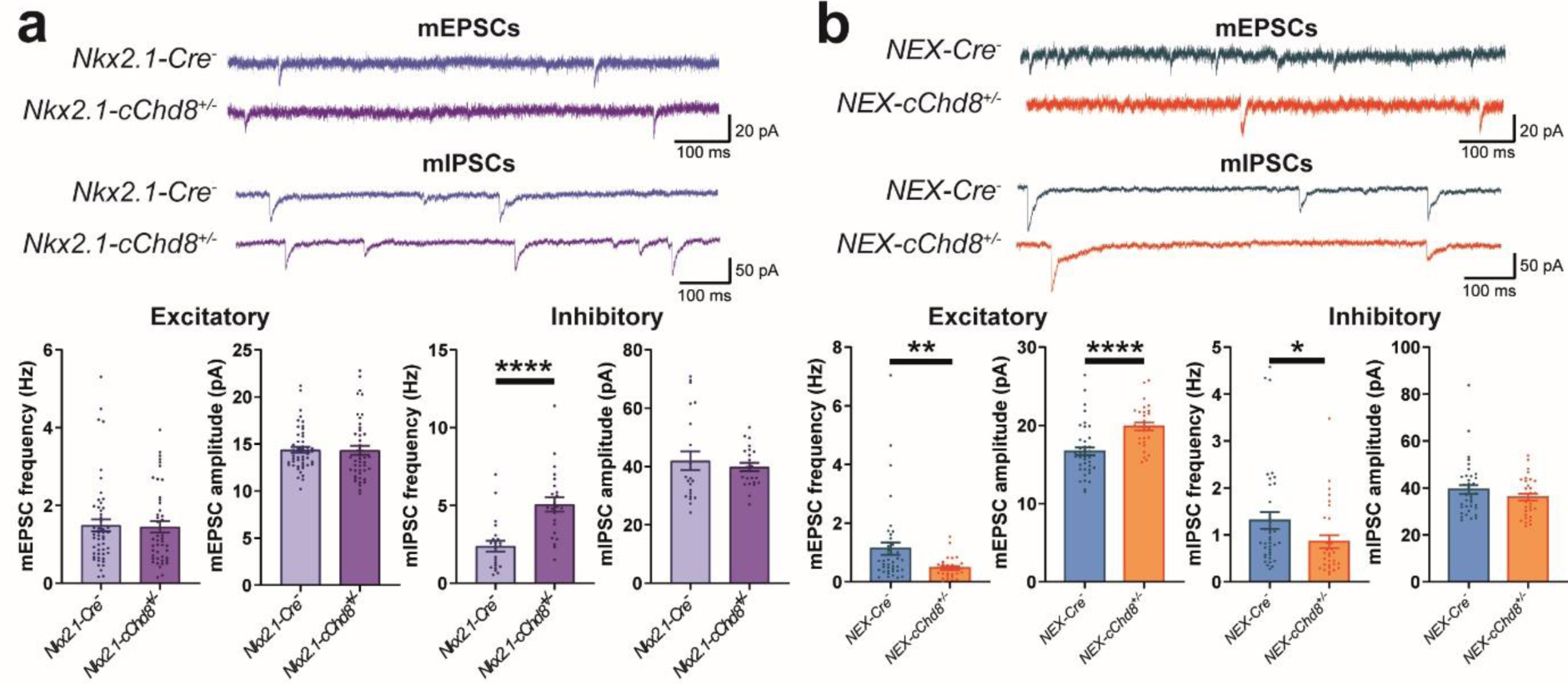
Differential alterations to synaptic transmission from cell-type-specific Chd8 heterozygosity. (**a, b**) Representative mEPSC (*top*) and mIPSC (*bottom*) recordings from P19-21 *Nkx2.1-cChd8*^*+/-*^ (**a**, *purple*) and *NEX-cChd8*^*+/-*^ (**b**, *orange*) neurons and respective controls (*blue and teal traces*) alongside quantifications of mEPSC and mIPSC frequency and amplitude. *Nkx2.1-cChd8*^*+/-*^ neurons showed no difference in mEPSC frequency or amplitude but showed increased mIPSC frequency (*p*<0.0001) with no change in amplitude. *NEX-cChd8*^*+/-*^ neurons showed decreased mEPSC frequency (*p*=0.0027) and increased mEPSC amplitude (*p*<0.0001) with decreased mIPSC frequency (*p*=0.011) and no change in mIPSC amplitude. *p<0.05, **p<0.01, ****p<0.0001

### Disrupted homeostatic regulation of synaptic transmission in Chd8^+/-^ neurons

To examine homeostatic responses in deep layer PFC neurons, we developed an *ex vivo* acute slice paradigm (Fig. 4a; see Materials and Methods for details). We found that a 6-hour incubation with TTX and APV to reduce network activity reliably caused an increase in mEPSC frequency in P13-15 WT PFC neurons, with no change in amplitude (Fig. 4b). When we tested *Chd8*^+*/-*^ PFC neurons with the same paradigm, they failed to respond (Fig. 4b). Interestingly, we were unable to elicit plasticity through treatment with TTX alone (Fig. 4b), suggesting a mechanism dependent on blocking NMDA receptors from stochastically released glutamate. We were also unable to elicit changes to inhibitory transmission with TTX alone (Fig. 4c), therefore we trialled a TTX plus Gabazine (GZ) treatment in order to block stochastic GABA transmission. This had no effect on WT neurons but triggered an abnormal response in *Chd8*^+*/-*^ neurons, resulting in a significant increase in mIPSC frequency with no change in amplitude (Fig. 4c). Together these results show that reduced *Chd8* expression can disrupt the regulation of homeostatic plasticity specifically mediated by spontaneous neurotransmission.

**Fig. 4:**
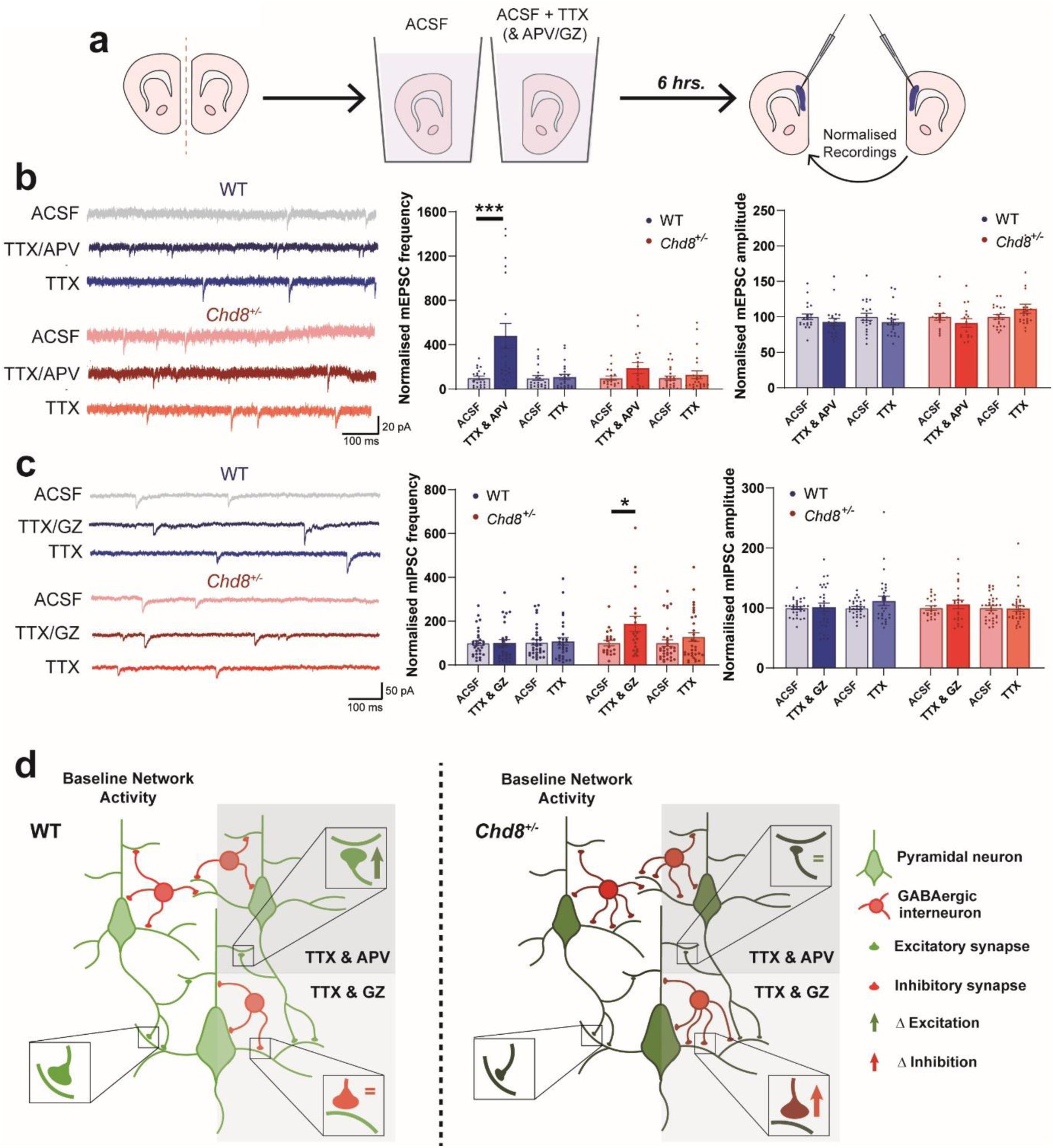
Disrupted homeostatic regulation of synaptic transmission in Chd8^+/-^ neurons. (**a**) Schematic of homeostatic plasticity induction paradigm in *ex vivo* P13-15 slices. (**b**) mEPSC responses to blocking AP firing +/- NMDA receptors; left: representative mEPSC recordings from WT (*blue*) and *Chd8*^*+/-*^ (*red*) neurons incubated for 6 hours in ACSF (*top*), ACSF+TTX+APV (*middle*) or ACSF+TTX (*bottom*) alongside quantifications of frequency (*middle*) and amplitude (*right*). WT neurons displayed a significant increase in mEPSC frequency following TTX/APV treatment (*p*=0.0003; genotype effect, *p*=0.029) while *Chd8*^*+/-*^ neurons failed to respond; no effects were seen on mEPSC amplitude. Treatment with TTX alone did not induce any change in mEPSC frequency or amplitude in either WT or *Chd8*^+*/-*^ neurons. (**c**) mIPSC responses to blocking AP firing +/- GABA_A_ receptors; *left:* representative mIPSC recordings arranged as in (**b**). WT neurons displayed no change in mIPSC frequency following TTX/GZ treatment while *Chd8*^+*/-*^ neurons showed a significant increase (*p*=0.021), with no effects on mIPSC amplitude. Treatment with TTX alone did not affect mIPSC frequency or amplitude in either WT or *Chd8*^+*/-*^ neurons. (**d**) Summary Fig. describing the synaptic phenotype resulting from *Chd8* haploinsufficiency. Reduced *Chd8* expression throughout development establishes a reduced E:I balance through decreased functional excitatory transmission (symbolized by smaller excitatory synapses) and increased numbers of inhibitory synapses onto basal dendrites. Altered homeostatic responses are seen in *Chd8*^+*/-*^ neurons. *p<0.05, ***p<0.001.

## Discussion

Overall, these results show that reduced expression of *Chd8* significantly alters synaptic development within the mouse PFC in a highly dynamic, stage specific manner that acts largely cell-autonomously in individual cell types to produce contrasting changes in excitatory and inhibitory synaptic transmission. Furthermore, *Chd8*^+*/-*^ neurons are unable to adequately retune excitatory synaptic transmission to respond to reductions in spontaneous neurotransmission, instead showing inappropriate increases in the level of inhibitory synaptic transmission.

Two previous studies have reported alterations to miniature postsynaptic currents in *Chd8*^+*/-*^ mouse models concurrent with altered social behaviours. In contrast to our findings, Platt *et al.* found no change in excitatory transmission within the nucleus accumbens of adult mice and reported reduced inhibitory transmission (8). Jung *et al.* examined the upper layers of the cortex and hippocampus of juvenile animals with a heterozygous, frame-shift point mutation of *Chd8*, finding no changes to synaptic transmission in cortex, and alterations specifically to inhibitory transmission in hippocampus (7). These discrepancies likely result from two key differences between our studies; the brain regions in which synaptic transmission was surveyed and the developmental stage of the animals. We have provided a thorough characterization of the impact of reduced *Chd8* expression on E:I balance within pyramidal neurons of the deep layers of the PFC, a cell type and brain region that are highly relevant to ASD etiology (34,35). Additionally we have gone beyond previous studies by examining synaptic transmission across neurodevelopment, showing that changes to E:I balance are highly dynamic and dependent on the age of the animals. Strikingly, our study has revealed a key developmental window during the adolescence (P14-P20) of *Chd8*^+*/-*^ mice where E:I balance is substantially reduced, characterized by decreased excitatory and increased inhibitory synaptic transmission, and outside of which synaptic transmission is relatively similar to WT animals. Such developmental stage-specific changes to synaptic transmission have been reported in a *Shank3B*^*-/-*^ mouse (11), which together with our work highlights that a longitudinal examination of synaptic transmission is necessary to fully understand how E:I balance is impacted at different critical time periods in animal models of ASD.

By utilizing conditional Cre drivers we have demonstrated that cell-type-specific reduction of *Chd8* expression is sufficient to alter excitatory and inhibitory synapses in these cell types, providing new insight into how reduced *Chd8*^+*/-*^ expression impacts synaptic development and function. Our findings indicate that synaptic transmission phenotypes do not emerge through a global reduction in *Chd8* expression across neural circuits but rather result from discrete mechanisms arising within individual neuronal subtypes. Specifically, changes to inhibitory transmission are driven entirely by the reduction of *Chd8* expression in interneurons, while for excitatory synapses the reduction in mEPSC frequency appears to be the core, cell-autonomous phenotype with other changes possibly resulting from a complex interplay of different cellular responses. It is also interesting to note that the impact of reduced *Chd8* expression on inhibitory synaptic changes is to a significant extent structural, affecting synaptic densities, while our data are suggestive of primarily functional effects on excitatory neurotransmission. Determining how these potentially distinct mechanisms operate in different neuronal subtypes will be an essential avenue for future research.

Within this study we have developed a novel and robust assay that utilizes global pharmacological blockade of action potential firing and spontaneous neurotransmission to induce homeostatic synaptic responses in *ex vivo* slices of PFC. Interestingly, blockade of action-potential firing alone was found to be insufficient to induce plasticity changes, suggesting a mechanism that is dependent upon the blockade of spontaneous neurotransmitter release. Rapid homeostatic scaling, although affecting mEPSC amplitude as opposed to frequency, has been previously described for the blockade of spontaneous glutamate transmission through NMDA receptors in cultured hippocampal neurons (43). We now also describe a novel finding of plasticity changes being initiated through the blockade of spontaneous GABA transmission. By utilizing this assay, we have shown, for the first time, that homeostatic plasticity mechanisms are highly dysregulated in the PFC of *Chd8*^+*/-*^ mice. Specifically, *Chd8*^+*/-*^ neurons display a blunted ability to induce compensatory changes in excitatory synaptic transmission and abnormal, inappropriate changes to inhibitory synaptic transmission. A recent study examining four separate ASD mouse models displaying altered E:I balance within the sensory cortex showed that these changes acted to stabilize synaptic depolarization and spiking and that the mice showed little to no change in sensory-evoked activity as a result (21). They therefore proposed that changes to E:I balance may be the end point of homeostatic compensatory mechanisms acting to stabilize circuit excitability rather than a causal mechanism of ASD (21). Our results are not consistent with this argument as they show that homeostatic plasticity mechanisms themselves can be profoundly dysregulated in ASD models. It should be noted that all the models examined in the study from Antoine *et al.* displayed increased E:I ratios, whereas we find that *Chd8*^+*/-*^ mice show a profoundly decreased E:I balance. Therefore, differential mechanisms may be acting to alter cortical circuitry, perhaps indicative of differential pathophysiological subgroups. Our findings are in agreement with a recent report showing that homeostatic plasticity mechanisms are defective in neuronal cultures with reduced *Shank3* expression and in primary visual cortex following eyelid suture in *Shank3* knockout mice (20). We now provide evidence that homeostatic responses are both insufficient and aberrant in an ASD model within neuronal circuitry more directly relevant to ASD pathology. Therefore, we propose that dysregulation of homeostatic plasticity contributes to the establishment of a reduced E:I balance in the PFC of *Chd8*^+*/-*^ mice (Fig. 4d), thereby providing a potential mechanism through which mutation of *CHD8* can impair the function of ASD-relevant circuits.

## Supporting information

Supplemental Methods and Statistics

## Acknowledgements

This work was funded by grants from the Simons Foundation (SFARI #344763 to M.A.B. and #653443 to M.A.B. and L.C.A.), the BBSRC (BB/P000479/1 to L.C.A) and a KHP Challenge Fund Award (R160801 to L.C.A). L.C.A was supported by a NARSAD Young Investigator Grant from the Brain & Behavior Research Foundation. R.A.E. was supported by the King’s Bioscience Institute and the Guy’s and St Thomas’ Charity Prize PhD program in Biomedical and Translational Science. The authors wish to thank Lorcan Browne, Nathalie Higgs, Ruth Taylor, Carly Schott and BSU staff for their technical assistance during the study. We thank the Marin and Rico laboratories for providing the Nkx2.1-Cre and NEX-Cre lines.

