## Supplemental Methods and Statistics for "Cell-type-specific synaptic imbalance and disrupted homeostatic plasticity in cortical circuits of ASD-associated *Chd8* haploinsufficient mice"

### Supplementary Materials and Methods

#### Mice and genotyping

Experimental *Chd8*$^{+/-}$ mice were produced by crossing *Chd8*$^{+/-}$ with C57BL/6J mice, ensuring equal paternal and maternal inheritance of the *Chd8*$^{-}$ allele. For all genotyping, genomic DNA was extracted from ear or tail samples. Primers used were:

*Chd8*$^{flox}$ primers (Forward (F) = GCC GAG GGG ATG AGG ATA TTT AGG, Reverse (R) = GGT ACA TAT GCC TTA AAA ATC AGG CCC AG) yield a WT band of 211bp and *Chd8*$^{flox}$ band of 276bp. *Chd8*$^{-}$ primers (F = CCC ACA TCA AGT GGC TGT AA, R = GGT AGG GAA GCA GTG TCC AG) yielded a WT band of 1.1kb and a *Chd8*$^{-}$ band of 395bp. *Cre* primers (F = CCT GGA AAA TGC TTC TGT CCG, R = CAG GGT GTT ATA AGC AAT CCC) yielded a *Cre* band of 390bp. *EGFP* primers (CCT ACG GCG TGC AGT GCT TCA GC, R = CGG CGA GCT GCA CGC TGC GTC CTC) yielded an *EGFP* band of approximately 350bp.

#### Electrophysiology

*Acute brain slice preparation*

Mice were anaesthetised by isofluorane inhalation and immediately decapitated. 300 µm thick anterior coronal slices containing the prefrontal cortex (PFC) were prepared from dissected brains using a VT1000S vibratome (Leica). Brains remained in ice-cold cutting solution (240 mM sucrose, 5 mM KCl, 1.25 mM $Na_2PO_4$, 2 mM $MgSO_4$, 1 mM $CaCl_2$, 26 mM $NaHCO_3$ and 10 mM D-glucose) equilibrated with 95% $O_2$/5% $CO_2$ throughout the dissection and slicing procedures. Slices were allowed to recover in room-temperature artificial cerebrospinal fluid (ACSF; 124 mM NaCl, 5 mM KCl, 1.25 mM $Na_2HPO_4$, 2 mM $MgSO_4$, 2 mM $CaCl_2$, 26 mM $NaHCO_3$ and 20 mM D-glucose) equilibrated with 95% $O_2$/5% $CO_2$ for one hour before recording. For adult mice (P55-P60) an N-methyl-D-glucamine (NMDG) cutting solution (92 mM NMDG, 92 mM HCl, 2.5 mM KCl, 1.2 mM Na $H_2PO_4$, 30 mM $NaHCO_3$, 20 mM HEPES, 25 mM glucose, 5 mM sodium ascorbate, 2 mM thiourea, 3 mM sodium pyruvate, 10 mM $MgSO_4$, 0.5 mM $CaCl_2$) was used followed by $Na^+$ reintroduction (41) to aid neuronal survival.

*Whole-cell patch clamp electrophysiology*

Whole-cell patch clamp recordings were taken from cortical pyramidal projection neurons located within layers V and VI of the PFC. Slices were continuously perfused with ACSF equilibrated with 95% $O_2$/5% $CO_2$, held in position using a platinum wire harp and visualised using an Olympus BX51WI microscope and Rolera Bolt camera under a 40x water-dipping objective. Neurons with a large diameter and pyramidal-shaped soma within the correct cortical region were targeted for recordings. This method was presumed to largely exclude inhibitory interneurons, which are known to have small, round somas and represent a minority of cells within the cortex (<20%). All recordings were performed at room temperature using a Patch clamp EPC 10 USB amplifier, Digidata 1440 digitiser and PatchMaster software (HEKA) with signals filtered at 10 kHz and sampled at 50 kHz. All traces were recorded and analyzed blind to genotype.

*Miniature postsynaptic current recordings*

For miniature postsynaptic current recordings 1 µM tetrodotoxin (TTX, Tocris) was added to the ACSF to prevent action potential firing. Additionally, either 10 µM SR-95531 (Gabazine, Tocris) or 10 µM 2,3-dihydroxy-6-nitro-7-sulfamoyl-benzo[f]quinoxaline (NBQX, Tocris) and 25 µM (2*R*)-amino-5-phosphonovaleric acid (D-APV, Tocris) were added to isolate mEPSCs or mIPSCs respectively. Borosilicate glass electrodes (3-5 MΩ resistance) were filled with K-gluconate internal solution (135 mM K-gluconate, 10 mM KCl, 10 mM HEPES, 1 mM $MgCl_2$, 2 mM Na-adenosine triphosphate ($Na_2ATP$) and 0.4 mM Na-guanosine triphosphate ($Na_3GTP$)) for mEPSC recordings while a $Cl^-$-loaded internal solution (150 mM CsCl, 1.5 mM $MgCl_2$, 0.5 mM EGTA, 10 mM HEPES, 4 mM $Na_2ATP$, 0.4 mM $Na_3GTP$) was used for mIPSC recordings. Three 60-second traces of spontaneous activity (consisting of 60x 1 second sweeps) were recorded for each neuron with membrane potential clamped at -70 mV from which the average mEPSC/mIPSC frequency and amplitude were determined. A square voltage-step pulse (± 10 mV for 10ms) was recorded before and after each trace to determine series resistance. Any cell determined to have series resistance values > 20 MΩ or whose series resistance varied by >20% over the course of recording, were excluded from further analysis. The resulting traces were analysed using MiniAnalysis Program 6.0.3 software (Synaptosoft).

*Intrinsic cell properties recordings*

All intrinsic cell properties were recorded in standard ACSF using K-gluconate internal solution. Upon cell break in, resting membrane potential (RMP) was recorded immediately to minimise the impact of dialysis with internal solution. RMP was determined by switching to current clamp mode and injecting 0 current then compensating for liquid junction potential (14.7 mV). In voltage clamp mode a square voltage-step pulse (± 10 mV for 10 ms) was then administered. Whole-cell capacitance was calculated as the integral of the resulting capacitive transient and membrane resistance (*Rm*) was calculated from the resulting current step according to Ohm's law (*Rm = V/I*). The frequency of action potential firing versus current stimulus (*f-I* curves) was then measured in current clamp using a 12-step 1 second current injection protocol starting at -80 pA and increasing in 50 pA intervals. To assess action potential characteristics, a 10-step protocol with a finer 5 pA interval was used in order to elicit voltage traces containing a single action potential. The amplitude, width and firing threshold of this single event were then determined using a custom-written MATLAB script (MathWorks).

**Sholl Analysis**

Mice were sacrificed at postnatal day 22 (P22) by rising $CO_2$ concentration. Brains were dissected and Golgi-Cox stained according to the FD Rapid GolgiStain Kit (FD Neurotechnologies) protocol. Before the stain development step, impregnated brains were embedded in 4% low-melting point agarose (Thermo Scientific) and cut into 100 µm coronal sections using a VT1000S vibratome (Leica). Brightfield Z-stacks of stained neurons were captured using an Eclipse Ti microscope (Nikon) under a 20x objective. Dendritic branching patterns were reconstructed in three dimensions from Z-stacks by manually tracing the basal dendritic tree of layer V/VI pyramidal projection neurons using the Image J plugin Simple Neurite Tracer(42). The in-built Sholl analysis function of Simple Neurite Tracer was used to perform the analysis with a radius step size of 5 microns.

**Synapse Analysis**

*Immunohistochemistry*

P14 and P20 *Chd8*$^{+/-}$*;Thy1-GFP-M* mice were deeply anaesthetised with Euthatal (Merial) and transcardially perfused with 5 ml phosphate buffered saline (PBS) followed by 5 ml 4% paraformaldehyde (PFA). Brains were dissected and postfixed overnight in 4% PFA before being embedded in 4% low-melting point agarose (Thermo Scientific) and cut into 100 µm coronal sections using a Leica VT 1000S vibratome. Sections containing the PFC were permeabilized in PBS + 1% Triton-X100 (PBS-T) for 4 hours then placed in block solution (3% Bovine Serum Albumin (BSA), 10% Foetal Bovine Serum (FBS), 0.2 M glycine, in PBS-T) overnight. Next, the sections were incubated at 4°C for 3 days in primary antibodies specific for green fluorescent protein (GFP; 1:1000, chicken anti-

GFP, Abcam) and vesicular GABA transporter (VGAT; 1:1000, rabbit anti-VGAT, Synaptic Systems) diluted in block solution. Sections were washed with PBS-T then incubated overnight at room temperature in secondary antibodies (goat α-chicken Fluor488 and goat α-rabbit Fluor568, 1:2000, Alexa) diluted in block solution. Sections were washed with PBS-T and PBS then mounted onto glass slides with Mowiol (Sigma).

*Image acquisition and analysis*

Z-stack images were taken of dendrites (and associated VGAT staining) from GFP+ layer V/VI projection neurons within the PFC using either a Zeiss LSM 800 or Nikon A1R Point-Scanning Confocal Microscope under 63x or 100x oil immersion objectives. A single secondary apical and basal dendrite were imaged per neuron and a minimum of 4 neurons were imaged per animal. The dendrites and the spines were reconstructed using the filament tracer tool within IMARIS software (BitPlane) from which dendrite length and spine density were quantified. The Spots function was then used to detect the VGAT staining puncta within 1µm of the dendritic shaft. The mean VGAT puncta size was calculated for each animal and used as the estimated XY diameter during spot detection so as to normalise synapse detection between animals. All imaging and analysis were performed blind to sample genotype.

### Supplementary Tables:

**Supplementary Table 1:** *Descriptive statistics and results of pairwise analyses*

| | Pairwise Analyses (Mann-Whitney U & t-test) | | | | | |
|---|---|---|---|---|---|---|
| **Comparison (Units)** | **Genotype** | **N (Neurons (animals))** | **Mean** | **S.E.M (±)** | **Test** | **P-value** |
| **P20 mEPSC frequency (Hz)** | WT | 38 (6) | 1.023 | 0.2 | MW-U | **0.0027** |
| | *Chd8*$^{+/-}$ | 44 (8) | 0.46 | 0.071 | | |
| **P20 mEPSC amplitude (pA)** | WT | 38(6) | 14.78 | 0.88 | MW-U | **< 0.0001** |
| | *Chd8*$^{+/-}$ | 45 (8) | 10.87 | 0.54 | | |
| **P20 mIPSC frequency (Hz)** | WT | 62 (8) | 3.97 | 0.28 | t-test | **0.0009** |
| | *Chd8*$^{+/-}$ | 55 (7) | 5.73 | 0.43 | | |
| **P20 mIPSC amplitude (pA)** | WT | 62 (8) | 43.71 | 1.011 | MW-U | **0.14** |
| | *Chd8*$^{+/-}$ | 55 (7) | 41.39 | 1.33 | | |
| **AP amplitude (mV)** | WT | 54 (8) | 59.33 | 1.25 | MW-U | **0.2605** |
| | *Chd8*$^{+/-}$ | 42 (6) | 61.44 | 0.52 | | |
| **AP width-at-half-height (ms)** | WT | 54 (8) | 1.89 | 0.034 | t-test | **0.4228** |
| | *Chd8*$^{+/-}$ | 42 (6) | 1.84 | 0.04 | | |
| **Rheobase (pA)** | WT | 54 (8) | 90.67 | 6.99 | MW-U | **0.473** |
| | *Chd8*$^{+/-}$ | 42 (6) | 85.36 | 7.54 | | |
| **AP firing threshold (mV)** | WT | 54 (8) | 34.38 | 0.62 | MW-U | **0.3814** |
| | *Chd8*$^{+/-}$ | 42 (6) | 34.77 | 0.79 | | |
| **Resting membrane potential (mV)** | WT | 55 (8) | -74.77 | 0.7 | MW-U | **0.7742** |
| | *Chd8*$^{+/-}$ | 46 (6) | -74.09 | 0.88 | | |
| **Whole-cell capacitance (pF)** | WT | 62 (8) | 157.6 | 6.62 | t-test | **0.2677** |
| | *Chd8*$^{+/-}$ | 51 (6) | 168.4 | 6.93 | | |
| **Membrane Resistance (MΩ)** | WT | 62 (8) | 250.4 | 12.8 | MW-U | **0.1505** |

| | | | | | | |
|---|---|---|---|---|---|---|
| | *Chd8*$^{+/-}$ | 51 (6) | 241 | 20.39 | | |
| **P5 mEPSC frequency (Hz)** | WT | 28 (7) | 0.082 | 0.016 | MW-U | **0.3508** |
| | *Chd8*$^{+/-}$ | 32 (4) | 0.12 | 0.028 | | |
| **P14 mEPSC frequency (Hz)** | WT | 34 (5) | 0.48 | 0.078 | MW-U | **0.4954** |
| | *Chd8*$^{+/-}$ | 38 (4) | 0.36 | 0.041 | | |
| **P55-60 mEPSC frequency (Hz)** | WT | 39 (6) | 1.75 | 0.29 | MW-U | **0.1224** |
| | *Chd8*$^{+/-}$ | 29 (5) | 1.072 | 0.25 | | |
| **P5 mEPSC amplitude (pA)** | WT | 27 (7) | 13.38 | 1.12 | MW-U | **0.7454** |
| | *Chd8*$^{+/-}$ | 32 (4) | 13.14 | 0.68 | | |
| **P14 mEPSC amplitude (pA)** | WT | 34 (5) | 14.84 | 1.002 | MW-U | **0.0004** |
| | *Chd8*$^{+/-}$ | 38 (4) | 11.26 | 0.5 | | |
| **P55-60 mEPSC amplitude (pA)** | WT | 39 (6) | 13.4 | 0.41 | MW-U | **0.5296** |
| | *Chd8*$^{+/-}$ | 29 (5) | 14.04 | 0.91 | | |
| **P5 mIPSC frequency (Hz)** | WT | 40 (5) | 0.033 | 0.0058 | MW-U | **0.345** |
| | *Chd8*$^{+/-}$ | 40 (7) | 0.021 | 0.02 | | |
| **P14 mIPSC frequency (Hz)** | WT | 41 (6) | 1.62 | 0.21 | MW-U | **0.6239** |
| | *Chd8*$^{+/-}$ | 38 (7) | 1.76 | 0.23 | | |
| **P55-60 mIPSC frequency (Hz)** | WT | 38 (6) | 6.67 | 0.79 | MW-U | **0.8018** |
| | *Chd8*$^{+/-}$ | 26 (5) | 6.43 | 0.91 | | |
| **P5 mIPSC amplitude (pA)** | WT | 34 (5) | 29.88 | 2.035 | MW-U | **<0.0001** |
| | *Chd8*$^{+/-}$ | 37 (7) | 18.4 | 1.14 | | |
| **P14 mIPSC amplitude (pA)** | WT | 41 (6) | 34.78 | 1.18 | t-test | **0.008** |
| | *Chd8*$^{+/-}$ | 38 (7) | 42.15 | 1.72 | | |
| **P55-60 mIPSC amplitude (pA)** | WT | 38 (6) | 27.15 | 1.23 | MW-U | **0.0328** |
| | *Chd8*$^{+/-}$ | 26 (5) | 32.36 | 1.955 | | |
| **P14 apical spines (per 10 µm)** | WT | 29 (5) | 7.12 | 0.44 | MW-U | **0.5595** |
| | *Chd8*$^{+/-}$ | 30 (6) | 7.074 | 0.44 | | |
| **P14 basal spines (per 10 µm)** | WT | 26 (5) | 5.054 | 0.44 | MW-U | **0.0613** |
| | *Chd8*$^{+/-}$ | 32 (6) | 6.122 | 0.35 | | |
| **P14 apical VGAT puncta (per 10 µm)** | WT | 29 (5) | 2.55 | 0.31 | MW-U | **0.2823** |
| | *Chd8*$^{+/-}$ | 30 (6) | 3.68 | 0.71 | | |
| **P14 basal VGAT puncta (per 10 µm)** | WT | 26 (5) | 2.72 | 0.31 | MW-U | **0.2608** |
| | *Chd8*$^{+/-}$ | 32 (6) | 2.7 | 0.58 | | |
| **P20 apical spines (per 10 µm)** | WT | 28 (5) | 8.16 | 0.47 | t-test | **0.2722** |
| | *Chd8*$^{+/-}$ | 24 (5) | 7.53 | 0.32 | | |
| **P20 basal spines (per 10 µm)** | WT | 27 (5) | 8.11 | 0.56 | t-test | **0.5184** |
| | *Chd8*$^{+/-}$ | 23 (5) | 7.68 | 0.36 | | |
| **P20 apical VGAT puncta (per 10 µm)** | WT | 25 (5) | 3.76 | 0.52 | MW-U | **>0.9999** |
| | *Chd8*$^{+/-}$ | 24 (5) | 3.94 | 0.65 | | |
| **P20 basal VGAT puncta (per 10 µm)** | WT | 24 (5) | 3.63 | 0.63 | MW-U | **0.0002** |
| | *Chd8*$^{+/-}$ | 23 (5) | 6.89 | 1.086 | | |
| ***Nkx2.1* mEPSC frequency (Hz)** | *Cre*$^{-}$ | 51 (6) | 1.49 | 0.15 | MW-U | **0.7547** |
| | *cChd8*$^{+/-}$ | 45 (5) | 1.45 | 0.15 | | |
| ***Nkx2.1* mEPSC amplitude (pA)** | *Cre*$^{-}$ | 51 (6) | 14.39 | 0.31 | MW-U | **0.2952** |
| | *cChd8*$^{+/-}$ | 45 (5) | 14.34 | 0.49 | | |
| ***Nkx2.1* mIPSC frequency (Hz)** | *Cre*$^{-}$ | 22 (3) | 2.38 | 0.35 | MW-U | **< 0.0001** |
| | *cChd8*$^{+/-}$ | 23 (3) | 5.069 | 0.46 | | |
| ***Nkx2.1* mIPSC amplitude (pA)** | *Cre*$^{-}$ | 22 (3) | 42 | 3.26 | MW-U | **0.5815** |
| | *cChd8*$^{+/-}$ | 23 (3) | 39.85 | 1.44 | | |
| ***NEX* mEPSC frequency (Hz)** | *Cre*$^{-}$ | 41 (7) | 1.13 | 0.21 | MW-U | **0.0027** |
| | *cChd8*$^{+/-}$ | 30 (4) | 0.47 | 0.064 | | |
| ***NEX* mEPSC amplitude (pA)** | *Cre*$^{-}$ | 41 (7) | 16.68 | 0.51 | MW-U | **< 0.0001** |
| | *cChd8*$^{+/-}$ | 30 (4) | 19.87 | 0.49 | | |
| ***NEX* mIPSC frequency (Hz)** | *Cre*$^{-}$ | 38 (6) | 1.31 | 0.18 | MW-U | **0.0108** |
| | *cChd8*$^{+/-}$ | 31 (4) | 0.86 | 0.14 | | |
| ***NEX* mIPSC amplitude (pA)** | *Cre*$^{-}$ | 38 (6) | 39.34 | 1.84 | MW-U | **0.2877** |
| | *cChd8*$^{+/-}$ | 31 (4) | 36.12 | 1.51 | | |

**Supplementary Table 2:** *Descriptive statistics and results of multivariate analyses*

**Multivariate Analyses (2-way ANOVA)**

| Comparison (Units) | Genotype | Treatment | N (Neurons (animals)) | Mean | S.E.M (±) | Variable | F | P-value | Tukey's post hoc P-value |
|---|---|---|---|---|---|---|---|---|---|
| **Sholl Analysis** | WT<br>*Chd8*$^{+/-}$ | N/A | 45 (4)<br>24 (3) | N/A | N/A | Genotype | (1, 1163) = 0.3897 | 0.5326 | N/A |
| ***f* – *I* curves** | WT<br>*Chd8*$^{+/-}$ | N/A | 54 (8)<br>42 (6) | N/A | N/A | Genotype | (1, 96) = 0.07494 | 0.7849 | N/A |
| **Plasticity mEPSC frequency (normalised)** | WT | ACSF<br>TTX & APV | 20 (3)<br>19 (3) | 100<br>479.9 | 17.6<br>112.5 | Genotype | (1, 68) = 4.961 | 0.0292 | 0.0003 |
| | *Chd8*$^{+/-}$ | ACSF<br>TTX & APV | 17 (3)<br>16 (3) | 100<br>189.9 | 19.08<br>50.59 | | | | 0.9272 |
| | WT | ACSF<br>TTX | 21 (4)<br>23 (4) | 100<br>109.1 | 22.37<br>23.18 | Treatment | (1, 85) = 0.5352 | 0.4665 | 0.9948 |
| | *Chd8*$^{+/-}$ | ACSF<br>TTX | 22 (4)<br>23 (4) | 100<br>129.2 | 19.66<br>35.47 | | | | 0.9946 |
| **Plasticity mEPSC amplitude (normalised)** | WT | ACSF<br>TTX & APV | 20 (3)<br>20 (3) | 100<br>92.71 | 4.13<br>5.06 | Treatment | (1, 68) = 2.62 | 0.1102 | 0.8431 |
| | *Chd8*$^{+/-}$ | ACSF<br>TTX & APV | 17 (3)<br>16 (3) | 100<br>91.39 | 4.7<br>5.9 | | | | 0.8183 |
| | WT | ACSF<br>TTX | 21 (4)<br>23 (4) | 100<br>92.65 | 4.9<br>4.3 | Treatment | (1, 84) = 0.1662 | 0.6845 | 0.7102 |
| | *Chd8*$^{+/-}$ | ACSF<br>TTX | 22 (4)<br>22 (4) | 100<br>111.3 | 3.3<br>6.5 | | | | 0.3591 |
| **Plasticity mIPSC frequency (normalised)** | WT | ACSF<br>TTX & GZ | 31 (6)<br>29 (6) | 100<br>100.6 | 11.2<br>15.5 | Genotype | (1, 98) = 5.206 | 0.0247 | > 0.9999 |
| | *Chd8*$^{+/-}$ | ACSF<br>TTX & GZ | 21 (4)<br>21 (4) | 100<br>187.1 | 12.26<br>34.3 | | | | 0.0206 |
| | WT | ACSF<br>TTX | 33 (6)<br>30 (6) | 100<br>107,6 | 12.26<br>16.8 | Treatment | (1, 1190 = 1.593 | 0.2094 | 0.9347 |
| | *Chd8*$^{+/-}$ | ACSF<br>TTX | 33 (6)<br>32 (6) | 100<br>127.4 | 15.1<br>19.9 | | | | 0.6201 |
| **Plasticity mIPSC amplitude (normalised)** | WT | ACSF<br>TTX & GZ | 31 (6)<br>29 (6) | 100<br>101.3 | 2.4<br>6.7 | Treatment | (1, 98) = 0.4849 | 0.4878 | 0.9973 |
| | *Chd8*$^{+/-}$ | ACSF<br>TTX & GZ | 21 (4)<br>21 (4) | 100<br>106 | 3.5<br>7.04 | | | | 0.8788 |
| | WT | ACSF<br>TTX | 30 (6)<br>28 (6) | 100<br>112 | 2.5<br>7.7 | Treatment | (1, 119) = 1.379 | 0.2426 | 0.3357 |
| | *Chd8*$^{+/-}$ | ACSF<br>TTX | 33 (6)<br>32 (6) | 100<br>99.5 | 3.6<br>4.8 | | | | 0.9998 |
